# Senescence-associated *Cdkn1a* (p21) is upregulated in rodent dorsal root ganglion neurons across lesion-based neuropathic pain models

**DOI:** 10.64898/2026.01.20.700573

**Authors:** Philipp N. Ostermann, Rebecca Dinnendahl, Jörg Isensee, Tim Hucho

## Abstract

Senescence of peripheral sensory neurons has recently been associated with increased nociceptive signalling and pain chronification. Whether this association is found across pain conditions and independent study settings remains to be investigated. In this study, we made use of publicly available whole-genome transcriptome data sets on rodent dorsal root ganglion (DRG) tissue to investigate pain-related peripheral sensory neuron senescence across pain models and independent study settings. We focused on *Cdkn1a* (p21) and *Cdkn2a* (p16) expression levels as two widely-used senescence markers, and to explore their potential role in peripheral sensory neuron senescence. We found that *Cdkn1a* but not *Cdkn2a* RNA is significantly increased across different axotomy- and lesion-based neuropathic pain models, but less consistent in other pain models including inflammatory pain. We observed a sex-dependent effect of *Cdkn1a* upregulation following nerve injury, with significantly increased *Cdkn1a* RNA levels in DRG cells from female but not male rats. Lastly, *Cdkn1a* RNA levels are increased among all DRG neuronal subtypes, seem to reach their maximum three to seven days following nerve injury, and afterwards go back to baseline. These data suggest that *Cdkn1a* upregulation in DRG neurons is a widespread response to nerve injury that is found across independent study settings.

**Summary:** Analysis of publicly available whole-genome transcriptome data sets shows that senescence-associated *Cdkn1a* but not *Cdkn2a* expression is upregulated in DRG neurons across neuropathic pain models.

## Introduction

Chronic pain is a global burden affecting more than 20% of adults worldwide [5; 19; 38]. Since current concepts for treatment of chronic pain often fail to provide satisfying pain relief for the majority of patients, or cause therapy-limiting adverse effects, there is an urgent need for alternative pain resolving strategies [4; 11; 27; 42]. To develop such alternative strategies, we must elucidate the cellular mechanisms that drive enhanced nociceptor activity.

Neuronal senescence through aging and injury has recently been suggested as a mechanism leading to increased activity in peripheral sensory neurons and to pain chronification [6; 12; 36]. Reducing the rates of senescent peripheral sensory neurons with senolytics successfully alleviated nocifensive behaviour in two rodent neuropathic pain models so far [12; 36]. Following spared nerve injury (SNI), ablation of senescent neurons with ABT263 (Navitoclax) gradually improved mechanical allodynia and weight bearing in mice [12]. In a different study, Navitoclax, Venetoclax, and a senolytic PROTAC Bcl-X_L_ degrader were found to reduce or even resolve chronic constriction injury (CCI)-mediated hyperalgesia and allodynia [36]. Corroborating cellular senescence as a core aspect of lasting pain and identifying cellular markers thereof could open the way to a new class of analgesic drugs soon.

Cellular senescence is an apoptosis-resistant state of irreversible cell cycle arrest induced by various stressors and associated with the development of a senescence-associated secretory phenotype (SASP) [7; 14]. It is involved in tissue repair and preventing cancer, but SASP-related secretion of pro-inflammatory, pro-apoptotic, and pro-fibrotic factors can disturb tissue function and turn nearby cells themselves senescent [7].

Post mitotic non-proliferating neurons have been found to adopt a senescence-like state (previously proposed as “neurescence” [18]) in response to stressful environments [14; 20]. Senescent neurons are metabolically active, but show altered excitability and transition into the potentially contagious state of SASP similar to non-neuronal cells [14; 18]. Senescence of central nervous system neurons is increasingly associated with multiple neurodegenerative disease such as Alzheimer’s and Parkinson’s [15; 28]. The role of senescence of peripheral sensory neurons in lasting pain states is just emerging.

Cyclin-dependent kinase inhibitors CDKN1A (p21) and CDKN2A (p16) have been reported as markers for cellular senescence, both of which play a role in cell state stability as well as in induction and maintenance of cellular senescence [14; 34]. Senescence is associated with increased transcription, i.e. elevated *CDKN1A* and *CDKN2A* RNA levels. However, adoption of neuronal senescence is a complex, time-dependent process that may affect distinct neuronal subtypes differently [18]. Hence, senescence markers may not behave the same in different neuronal subtypes, and they are poorly defined for peripheral sensory neurons.

Proof-of-principle for a role of peripheral sensory neuron senescence for pain has been provided [12; 36]. The extent to which peripheral sensory neuron senescence occurs across different pain conditions, and different sensory neuron subtypes remains unclear. Moreover, the reliability of detecting increased senescence in peripheral sensory neurons under varying experimental conditions, using markers such as *Cdkn1a* and *Cdkn2a* gene expression, requires investigation.

Therefore, we made use of publicly available whole-genome microarray, bulk- and single-cell (sc)RNA sequencing data sets from previous studies on rodent dorsal root ganglion (DRG) tissue to investigate pain-related peripheral sensory neuron senescence across different, independent study settings. Such an approach is of high relevance due to the larger number of samples, the diversity of pain models and moreover, the independence of the performed studies. We hereby focused on *Cdkn1a* and *Cdkn2a* expression levels to investigate their potential as markers for peripheral sensory neuron senescence.

## Methods

### Sex and gender

Sex was considered a biological factor. Publicly available studies were analysed that used male, female, or male and female animals. We specifically addressed and discussed a potential role of sex in our study based on the available data sets.

### Use of publicly available data sets

We used publicly available transcriptomic data from studies listed in Table 1. Data from Uttam et al. were retrieved from supplementary tables 2 and 4 [39], from Parisien et al. from supplementary tables 1, 5, and 6 [25], from Strong et al. from supplementary table 2 [33], from Hu et al. from supplementary table 1 and 3 [17], from Renthal et al. from supplementary table 3 [26], from Cobos et al. from supplementary table 1 and the Gene Expression Omnibus (GEO) database (GSE102937) [10], from Starobova et al. from supplementary table S1 [30], from Athie et al. from supplementary table 1 [2], from Calls et al. from supplementary table 2 [6]. Data from Kummer et al. were downloaded from GEO (GSE104625) [23], from Vega-Avelaira et al. from GEO (GSE15041) [40], and data from Stephens et al. were provided by the authors upon request and read counts were downloaded from GEO (GSE100122) [32].

**Table 1.**
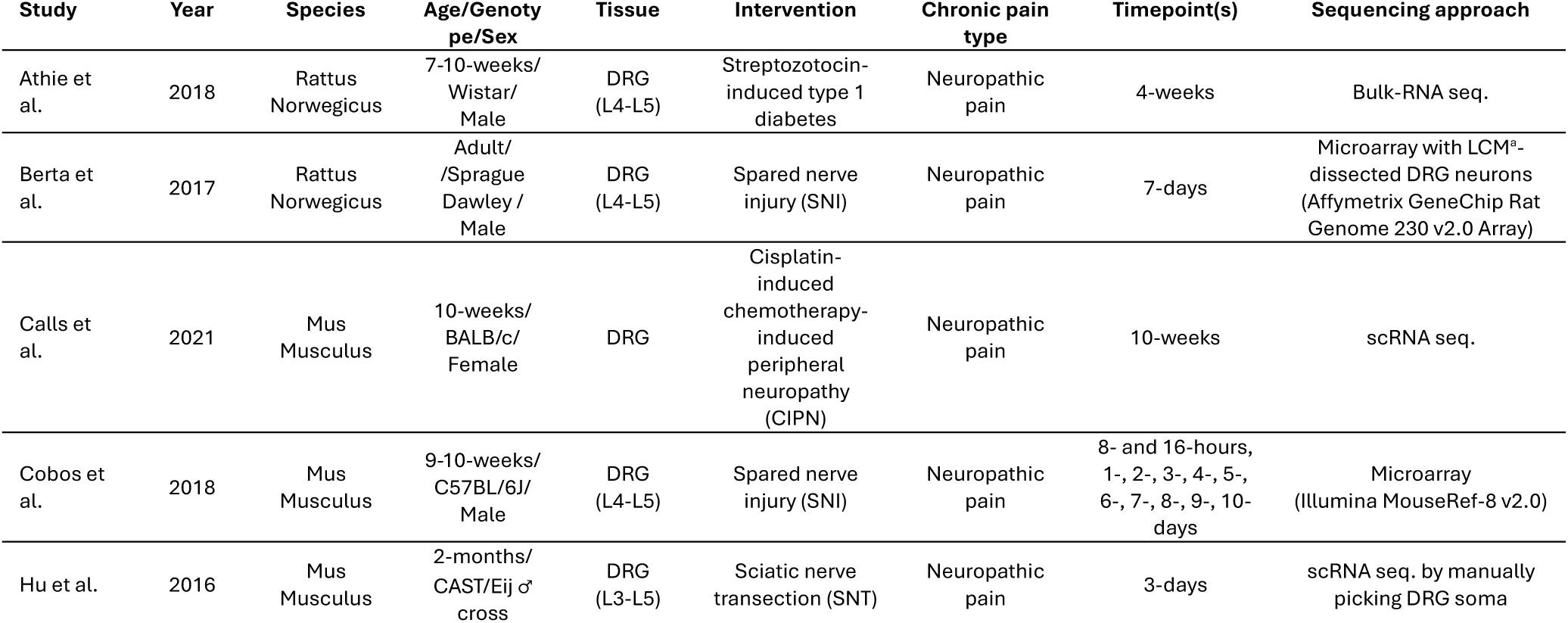
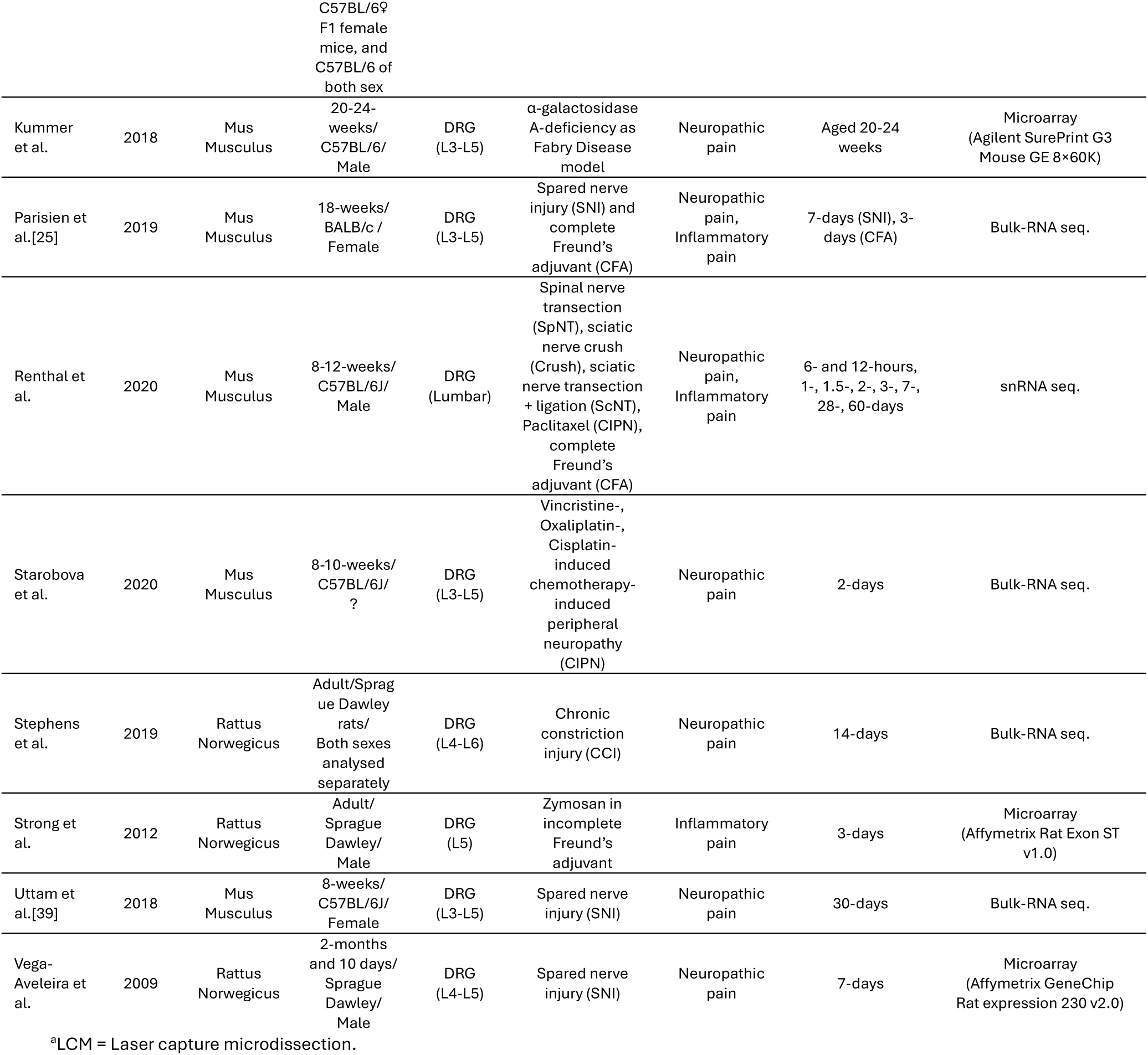
Microarray, bulk-RNA, and single-cell/nucleus RNA sequencing studies analysed.

### Software

Data were re-analysed using the online interactive tool GEO2R from the NCBI Gene Expression Omnibus (GEO) database [9; 13] utilizing the Bioconductor/R *limma* package [29], if processed data were not available as part of the original works (see below). GraphPad Prism (version 6) and RStudio with the *ggplot2* package [41] have been used for visualization and analysis of data. Inkscape (version 1.2) has been used to create figures. BioRender (Licence: University of Cologne – Medical School) has been used to create schematic illustrations.

### Statistical analysis

Data from Parisien et al. were processed in the original work using cuffdiff. Statistical significance has been determined as p < 0.05, |π| ≥ 1.11 as previously published [43]. Data from Strong et al. were processed in the original work with default parameters of the Genespring software to determine statistical significance (p-adj. < 0.05) following Benjamini-Hochberg correction for multiple testing. Data from Hu et al. were processed in the original work. Statistical significance for bulk-RNA sequencing was tested using the Wald test in DESeq2 and corrected for false discovery rates (FDR) using the Benjamini-Hochberg [17]. Statistical significance for scRNA-sequencing was determined using a Bayesian approach [22]. Data from Uttam et al. were processed in the original work following in-house R scripts as explained in Silva Amorim et al. [1], and by omitting genes with < 128 reads. Statistical significances were corrected for FDR using the Benjamini-Hochberg method [39]. Data from Stephens et al. were processed in the original work. Statistical significance was tested using the Wald test in DESeq2 and corrected for FDR using the Benjamini-Hochberg [32]. Data from Renthal et al. were processed in the original work. Statistical significance was tested using the QL F-test in edgeR [26]. Data from Cobos et al. were processed in the original work. Statistical significance was tested using a moderated F-statistic. Data from Vega-Avelaira et al. were re-analysed using the online interactive tool GEO2R from the GEO database [9; 13]. The analysis utilized the Bioconductor/R *limma* package [29] with default settings, including log-transformation of the data and FDR adjustment for multiple testing. Data from Starobova were processed in the original work. Statistical significance was tested using the Wald test in DESeq2 and corrected for FDR using the Benjamini-Hochberg [30]. Data from Athie et al. were processed in the original work. Statistical significance was tested using the Wald test in cuffdiff and corrected for FDR using the Benjamini-Hochberg [2]. Data from Kummer et al. were re-analysed using GEO2R [9; 13]. The analysis utilized the Bioconductor/R *limma* package [29] with default settings, including log-transformation of the data and FDR adjustment for multiple testing. Data from Calls et al. were processed in the original work. Statistical significance was tested using the default Seurat pipeline in R and corrected for FDR [6]. Statistical significance in the data set of Berta et al. [3] was tested by one-way ANOVA with Bonferroni multiple comparison testing between each condition (*p < 0.05). All significance thresholds are indicated throughout the main text.

## Results

### *Cdkn1a* RNA is consistently increased in dorsal root ganglion tissue across rodent lesion-based neuropathic pain models

It has recently been observed that inducing neuropathic pain in rodents leads to increased *Cdkn1a* (p21) and *Cdkn2a* (p16) RNA levels in DRG neurons, accompanied by an increased number of *Cdkn1a*- and *Cdkn2a*-positive DRG neurons [12]. First, we sought to test whether this increase in *Cdkn1a* and *Cdkn2a* RNA as a sign of cellular senescence is found across pain models and independent study settings. To this end, we made use of publicly available transcriptome data sets (Fig. 1).

**Fig. 1.**
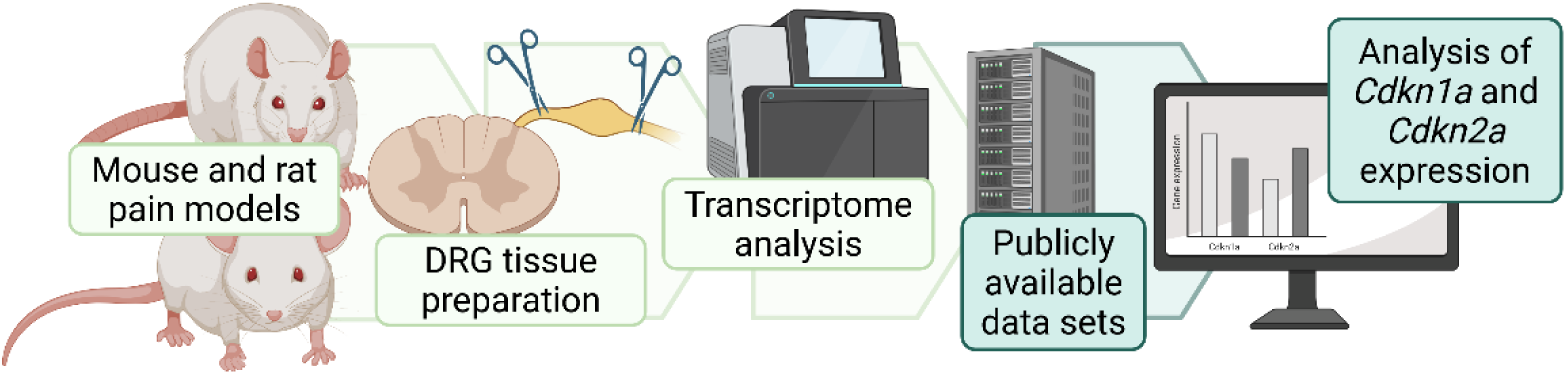
*Cdkn1a* (p21) and *Cdkn2a* (p16) expression analysis in dorsal root ganglion cells across rodent pain models. Schematic illustrating our study design with the here performed workflow in dark green, which builds on previously conducted transcriptome analyses of dorsal root ganglion (DRG) cell gene expression in mouse or rat pain models (light green).

We analysed 13 independent studies encompassing a total of 20 data sets of differential gene expression between rodent DRG-derived cells derived from pain models and respective controls (Table 1). Pain models included neuropathic pain models such as the SNI model or chemotherapy-induced peripheral neuropathy (CIPN), and inflammatory pain models based e.g., on complete Freund’s adjuvant (CFA). *Cdkn1a* and *Cdkn2a* RNA levels were analysed by whole-genome microarray, bulk-RNA, single-cell (sc), or single-nucleus (sn) RNA sequencing methods. Timepoints of transcriptome analysis ranged from 6 hours up to 70 days post-injury. For our analysis, we relied on the independent statistical analyses performed in the original works to obtain an unbiased picture of *Cdkn1a* and *Cdkn2a* differential gene expression levels across pain models, timepoints, and studies.

We found that *Cdkn1a* RNA is significantly increased in dorsal root ganglion tissue across rodent lesion-based neuropathic pain models (Fig. 2). In contrast, *Cdkn2a* RNA is rarely significantly increased in DRG tissue. *Cdkn2a* RNA was only found to be significantly increased 30 days following SNI [39], and 60 days following sciatic nerve transection (ScNT) [26]. Interestingly, *Cdkn1a* RNA was less consistently increased in neuropathic pain models that do not rely on direct lesioning of nerve tissue such as CIPN, streptozotocin-induced type 1 diabetes and a genetic Fabry disease model, although all these models supposedly model neuropathic pain mechanisms (Fig. 2). In the context of inflammatory pain, *Cdkn1a* RNA was only significantly increased in one out of three inflammatory pain data sets (Fig. 2).

**Fig. 2.**
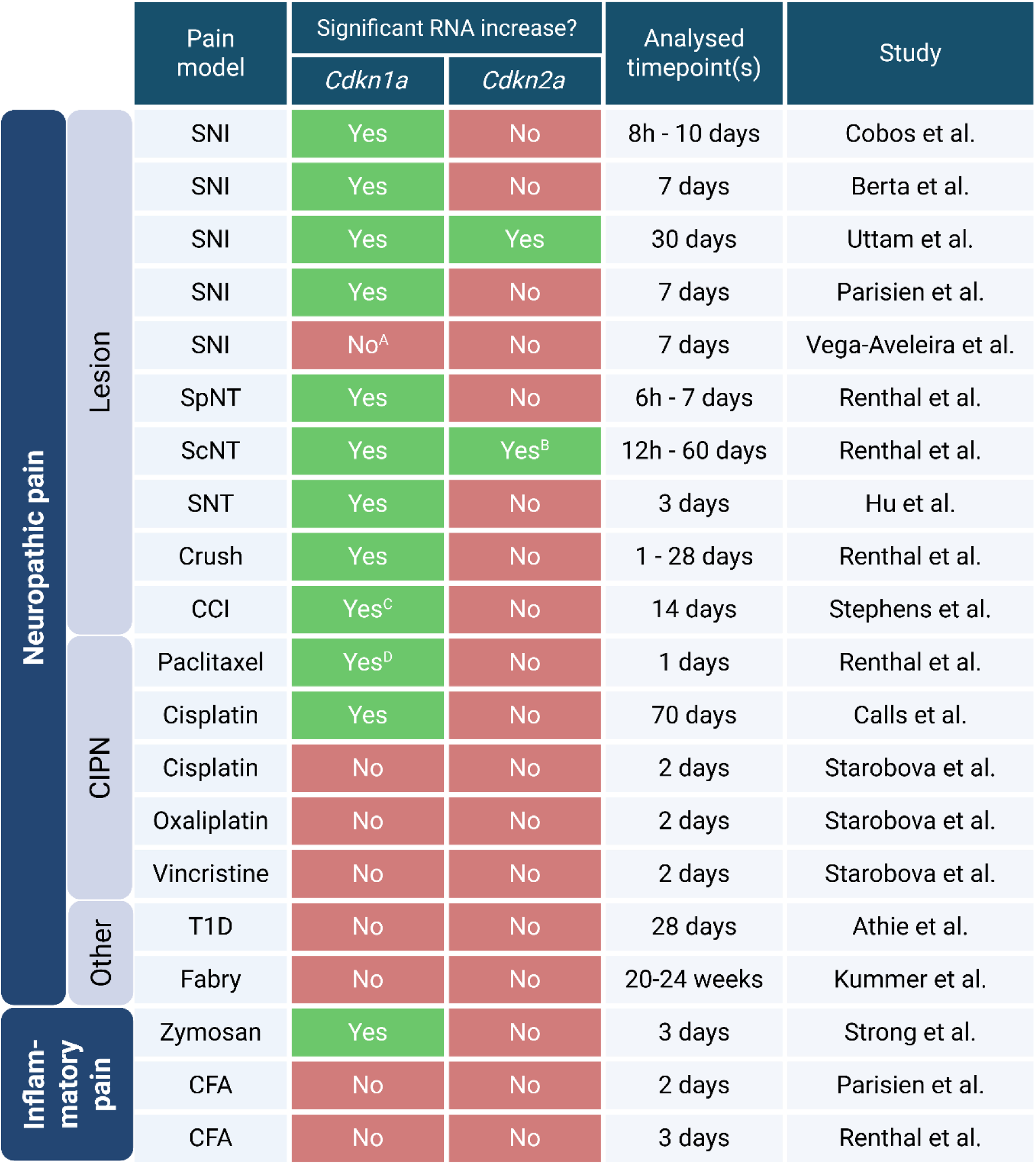
*Cdkn1a* (p21) RNA is significantly increased in dorsal root ganglion tissue across rodent lesion-based neuropathic pain models. CIPN = Chemotherapy-induced peripheral neuropathy; SNI = Spared nerve injury; SpNT = Spinal nerve transection; ScNT = Sciatic nerve transection and ligation; Crush = sciatic nerve crush; CCI = Chronic constriction injury; T1D = Streptozotocin-induced type 1 diabetes; Fabry = α-galactosidase A-deficiency as Fabry Disease model; CFA = Complete Freund’s adjuvant. ^A^not significant but p-adj. = 0.051 and Log2FC = 2.88 with n =3 samples. ^B^only at 60 days post-ScNT. ^C^only in DRG tissue from female but not male rats. ^D^only in *Sst*^+^ pruriceptors but not in other DRG neuron subtypes.

### *Cdkn1a* RNA is consistently increased in rodent DRG tissue after spared nerve injury

The most comprehensive work on peripheral sensory neuron senescence to date made use of the SNI model to study DRG neuron senescence and to test a potential pain resolving effect of senolytics [12]. To validate their findings, we now specifically investigated *Cdkn1a* and *Cdkn2a* RNA levels in independent studies that likewise used the SNI model.

In 2018, Cobos et al. performed whole-genome microarray analysis with ipsilateral DRG tissue isolated from mice at multiple timepoints over the course of ten days following SNI, and naïve (control) mice [10]. The onset of cold and tactile allodynia within five days, and which lasted for at least 15 days following SNI, was shown by acetone application and von Frey behavioural testing [10]. A total of 1,704 microarray probes were found to be differentially regulated over the course of ten days post-SNI when compared to the expression in DRG tissue from naïve control mice (p < 0.01) [10]. Three probes were directed against distinct parts of *Cdkn1a* mRNA allowing to exclude non-specific signals or technical errors, and all three probes showed significantly regulated RNA levels within the first ten days following SNI (F = 9.009, p < 0.00001, F = 26.96, p < 0.00001, F = 19.64, p < 0.00001) (Fig. 3A-C). The individual expression values indicated a trend towards early *Cdkn1a* upregulation within one day and a potential decline in *Cdkn1a* RNA from seven to eight days post-SNI on. In contrast, *Cdkn2a* RNA was not significantly regulated within these ten days [10].

**Fig. 3.**
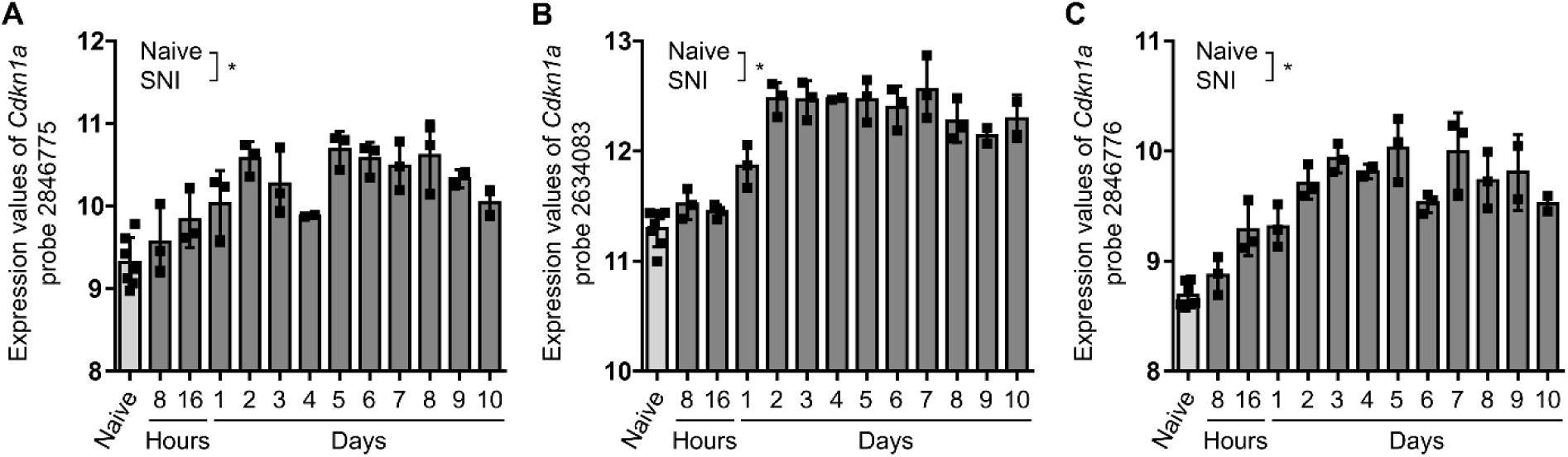
*Cdkn1a* (p21) RNA is increased in dorsal root ganglion tissue over ten days following spinal nerve injury in Cobos et al. (**A-C**) Expression values of the three *Cdkn1a*-specific probes 2846775 (A), 2634083 (B), and 2846776 (C) over 10 days following spinal nerve injury (SNI) obtained from Cobos et al. [10]. Statistical significance tested using moderated F-statistic in the original study (*p < 0.01): (A): F = 9.009, p < 0.00001; (B): F = 26.96, p < 0.00001; (C): F = 19.64, p < 0.00001 [10]. Data presented as individual data points and mean ± SD.

Parisien et al. also analysed gene expression changes following SNI [25]. The authors performed whole-genome bulk-RNA sequencing with lumbar DRG tissue from mice seven days after bilateral SNI to model neuropathic pain [25]. This timepoint was selected because von Frey behavioural testing showed that induced hypersensitivity peaked at seven days post-SNI [25]. In DRG tissue at seven days post-SNI, 1693 genes were significantly upregulated (p < 0.05, |π| ≥ 1.11) [25]. *Cdkn1a* but not *Cdkn2a* RNA was significantly increased at seven days post-SNI by 2.53-fold (p < 0.00005, |π| = 20.735).

Berta et al. focused on gene expression changes seven days post-SNI as well [3]. They analysed RNA levels specifically in injured and non-injured rat DRG neurons [3]. For this, the authors combined microarray analysis with tracer-guided laser-capture microdissection [3]. *Cdkn1a* expression was 5.77-fold higher in injured DRG neurons when compared to DRG neurons isolated from sham animals (Fig. 4A). *Cdkn1a* expression was unaffected in non-injured DRG neurons (Fig. 4A) [3].

**Fig. 4.**
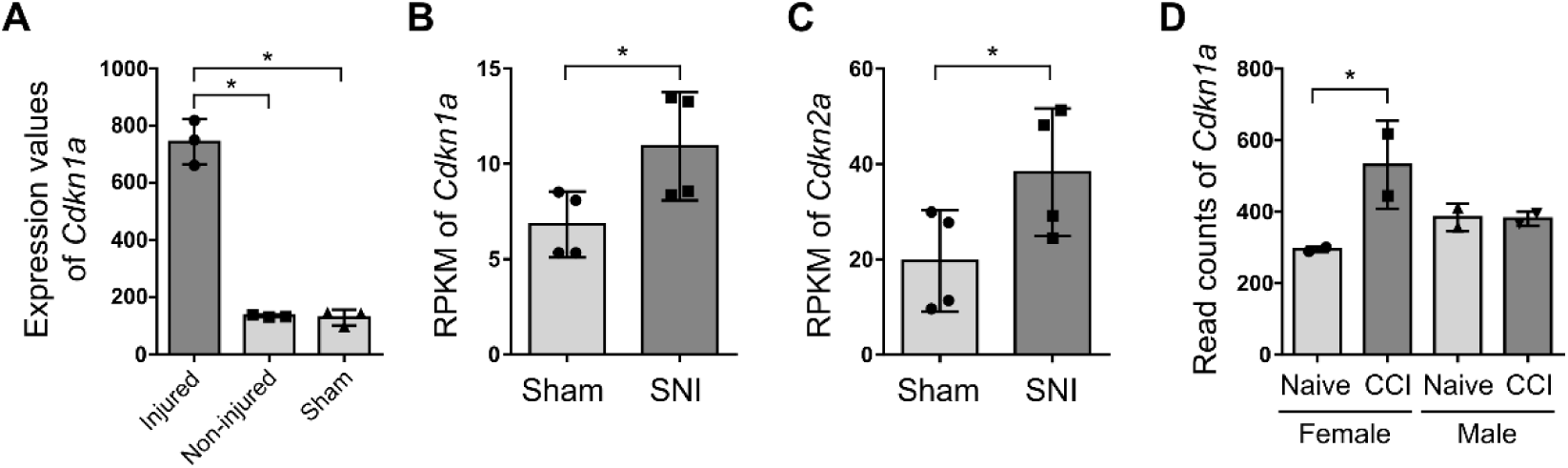
*Cdkn1a* (p21) RNA is increased in dorsal root ganglion tissue across rodent lesion-based neuropathic pain models. (**A**) Expression values of *Cdkn1a* RNA in injured and non-injured DRG neurons and DRG neurons from sham-operated animals seven days post-spinal nerve injury (SNI), obtained from Berta et al. [3]. Statistical significance tested by one-way ANOVA with Bonferroni multiple comparison testing between each condition (*p < 0.05). (**B, C**) Reads Per Kilobase of transcript per Million mapped reads (RPKM) of *Cdkn1a* (B) and *Cdkn2a* (C) RNA in DRG tissue 30 days post-SNI, obtained from Uttam et al. [39]. Statistical significance tested in the original study and corrected for false discovery rates (FDR) using the Benjamini-Hochberg method (*p-adj. < 0.05): (B): p-adj. = 0.0170; (C): p-adj. = 0.0030 [39]. (**D**) Read counts of *Cdkn1a* RNA in DRG tissue 14 days after chronic constriction injury (CCI), obtained from Stephens et al. [32]. Statistical significance tested in the original study using Wald test and corrected for false discovery rates (FDR) using the Benjamini-Hochberg (*p-adj. < 0.05) [32]. (**A-D**) Data presented as individual data points and mean ± SD.

A fourth study that investigated gene expression changes seven days post-SNI was performed by Vega-Aveleira et al. [40]. Microarray analysis was conducted with ipsilateral and contralateral DRG tissue from rats seven days after SNI or sham procedure [40]. The analysis was performed with two months- and ten days-old rats, and resulted in 206, and three significantly regulated probes, respectively (p < 0.05). In this microarray data, neither *Cdkn1a* nor *Cdkn2a* expression was found to be significantly regulated following SNI [40]. However, re-analysing RNA levels in ipsilateral and contralateral DRG tissue showed a trend towards increased *Cdkn1a* RNA levels with a log2-fold change of 2.88 and p-adj. of 0.051 (n = 3 samples) in two-months old but not ten-days old rats seven days post-SNI. Interestingly, a putative age-dependent effect on *Cdkn1a* upregulation and induced senescence in DRG neurons has been previously also observed when comparing 10-16 weeks-old with 20-24-months old mice following SNI [12].

In a fifth independent study using the SNI model, Uttam et al. performed whole-genome bulk-RNA sequencing with lumbar DRG tissue isolated from mice - this time 30 days - following bilateral SNI and mice that underwent sham surgery as control [39]. Von Frey behavioural testing confirmed significantly reduced mechanical thresholds at 14, 21, and 30 days following SNI as indication of chronic pain [39]. In their study, 144 mRNAs were found to be significantly increased 30 days following SNI (log2-fold change > 0.5, p-adj. < 0.05) [39]. Among these 144 mRNAs were *Cdkn1a* and *Cdkn2a* [39]. *Cdkn1a* RNA levels were significantly increased by 1.61-fold in DRG tissue 30 days following SNI (p-adj. = 0.0170), and *Cdkn2a* RNA levels were significantly increased by 1.72-fold (p-adj. = 0.0030) (Fig. 4B, C).

Overall, significantly increased *Cdkn1a* RNA levels in mouse and rat DRG tissue following SNI is a consistent finding across independent study settings, which therefore underlines the SNI model’s capacity to model induced *Cdkn1a* expression and potentially peripheral sensory neuron senescence in the context of lasting pain.

### *Cdkn1a* RNA is not consistently increased across neuropathic pain models of chemotherapy-induced peripheral neuropathy

In addition to surgical procedures, neuropathic pain can be modelled by CIPN using different drugs including cisplatin, paclitaxel, vincristine, and oxaliplatin. In 2021, Calls et al. performed scRNA sequencing analysis with DRG cells isolated from mice ten weeks after weekly administration of cisplatin or saline solution [6]. Onset of mechanical allodynia after three weeks was shown by von Frey behavioural testing [6]. Differential gene expression analysis between DRG-derived neurons after cell type clustering identified 122 significantly differentially expressed genes (p < 0.05) with *Cdkn1a* being the only gene with a p-adj. < 0.01 after correcting for false discovery rates. *Cdkn1a* RNA was increased by 1.99-fold between DRG neurons from cisplatin-treated when compared to control mice (p-adj. = 0.0018). Increased *Cdkn1a* expression in DRG neurons was confirmed on protein level (p21) by western blot analysis and immunocytochemistry [6]. Notably, the authors of this study analysed additional senescence-associated markers such as β-galactosidase activity and pH2AX, and to our knowledge were the first to suggest that cisplatin may induce neuropathic pain through a DRG neuronal senescence-like response [6]. *Cdkn2a* RNA levels were not significantly altered in the cisplatin model.

Starobova et al. performed whole-genome bulk-RNA sequencing analysis with DRG tissue from mice two days after vincristine, oxaliplatin, or cisplatin injection, and control mice as three different models for CIPN [30]. Vincristine led to 368, oxaliplatin to 295, cisplatin to 256 significantly differentially expressed genes (p-adj. < 0.05) [30]. *Cdkn1a* and *Cdkn2a* were not significantly differentially expressed in this study, which performed analyses at two days following compound injection. Given that Calls et al. observed significantly increased *Cdkn1a* expression 10 weeks following continuous cisplatin treatment [6], the timepoint of analysis may play an important role when analysing CIPN-induced *Cdkn1a* expression.

### *Cdkn1a* RNA is significantly increased only in DRG tissue from female but not male rats 14 days after chronic constriction injury

Previously, female sex has been associated with greater likelihood of senescence onset, although specific evidence is lacking (reviewed in [24]). Sex-specific differences in DRG neuron senescence have not been described.

Hence, we searched for prior sex-specific analyses of DRG neuron gene expression in the context of neuropathic pain to analyse *Cdkn1a* and *Cdkn2a* RNA expression. In 2019, Stephens et al. performed sex-specific whole-genome bulk-RNA sequencing analysis with DRG tissue from male and female rats 14 days after chronic constriction injury (CCI) and respective male and female untreated (naïve) controls [32]. Mechanical hypersensitivity at day 14 post-CCI was confirmed by von Frey behavioural testing [32].

The authors observed that 1185 genes were significantly upregulated in males but not in females, and that 146 gene were significantly upregulated in females but not in males upon CCI (log2FC > 0.5, p-adj. < 0.05). Among these 146 female-specific upregulated genes was *Cdkn1a*. *Cdkn1a* RNA levels were significantly increased by 2.06 -fold (p-adj. < 0.0001) in female rats 14 days post-CCI when compared to female naïve controls, but not significantly increased in male rats when compared to male naïve controls (Fig. 4D). This bulk-RNA sequencing data set was validated by *Cdkn1a*-specific RT-qPCR with a separate set of DRG tissue preparations. Interestingly, there was no significant sex-specific difference in *Cdkn1a* RNA levels found in DRG tissue derived from naïve rats [32]. This indicates a sex-specific response to nerve injury with no difference at baseline. *Cdkn2a* was not significantly differentially expressed at 14 following CCI. Hence, analysing the data by Stephens et al. indicated that *Cdkn1a* expression is differentially regulated in DRG neurons in male and female rats.

### *Cdkn1a* RNA is increased in all mouse DRG neuronal subtypes and to a lesser extent in non-neuronal cells across neuropathic pain conditions

So far, we have mostly analysed data from microarray or bulk-RNA sequencing studies that do not allow differentiating between *Cdkn1a* or *Cdkn2a* RNA in DRG-resident neuronal subtypes or associated non-neuronal cells. To now investigate whether *Cdkn1a* or *Cdkn2a* expression may predominantly occur in certain DRG neuron subtypes or additionally in non-neuronal cells, we searched for studies that performed single-cell analysis with neuronal subtype-specific downstream analysis.

In 2016, Hu et al. performed an initial whole-genome bulk-RNA sequencing analysis with pooled neurons from single-cell preparation derived from ipsilateral and contralateral DRG tissue isolated from mice three days after sciatic nerve transection (SNT) as control for their subsequent scRNA analysis [17]. This initial bulk-RNA sequencing experiment showed that 1179 genes were differentially expressed at three days post-SNT in the ipsilateral DRG tissue when compared to the contralateral (control) DRG tissue (fold change > 2, p-adj. < 0.05). Among these, 624 genes were upregulated including *Cdkn1a*. *Cdkn1a* RNA was significantly increased by 18.89-fold in this bulk-RNA analysis of pooled DRG neurons, already showing that *Cdkn1a* expression is indeed induced in DRG neurons (p-adj. = 0.0041).

Afterwards, the authors performed scRNA sequencing analysis based on the same experimental setting of isolating DRG tissue three days post-SNT. This way, differential gene expression was analysed individually in peptidergic nociceptors (222 up-, 181 down-regulated), in non-peptidergic nociceptors (1188 up-, 1067 down-regulated), and in large myelinated sensory neurons (51 up-, 32 down-regulated). *Cdkn1a* RNA was significantly increased in all three major sensory neuron populations at 3 days post-SNT: Peptidergic nociceptors = 3.86 log2-fold change, p-adj. < 0.00005; non-peptidergic nociceptors = 4.34 log2-fold change, p-adj. < 0.00005; large myelinated sensory neurons = 5.38 log2-fold change, p-adj. < 0.00005.

In the most comprehensive scRNA analysis in our context, Renthal et al. performed differential gene expression analysis with nine DRG neuronal subtypes and with DRG-resident non-neuronal cells. The authors performed snRNA sequencing analysis with DRG tissue across several pain models to map gene expression profiles in response to lesion-based nerve injury and CIPN [26]. For this, lumbar DRG tissue was isolated from adult mice following spinal nerve transection (SpNT), sciatic nerve crush (Crush), sciatic nerve transection and ligation (ScNT), and paclitaxel at various timepoints. Differential gene expression (p-adj. < 0.01 and log2fc > |1|) was captured between DRG cells from naïve (untreated) mice and their respective pain models within *Fam19a4*^+^/*Th*^+^ C-fiber low-threshold mechanoreceptor (cLTMR1) neurons, putative *Fam19a4*^+^/*Th*^low^ cLTMR2 neurons, *Mrgprd*^+^ non-peptidergic nociceptors (NPs), *Pvalb*^+^ proprioceptors (NF1, NF2), *Cadps2*^+^ Ad-LTMR (NF3) neurons, *Tac1*^+^/*Gpx3*^+^peptidergic nociceptors (PEP1), *Tac1*^+^/*Hpca*^+^ peptidergic nociceptors (PEP2), and *Sst*^+^ pruriceptors (SST) [26]. Furthermore, differential gene expression within DRG-resident myelinating Schwann cells, non-myelinating (Remark) Schwann cells, macrophages, endothelial cells, and fibroblasts was analysed [26].

*Cdkn1a* expression was significantly increased in all analysed DRG neuron subtypes across the three lesion-based neuropathic pain models SpNT (Fig. 5A), Crush (Fig. 5B), and ScNT (Fig. 5C). This increase occurred as early as 6 hours after surgical intervention in some DRG neuron subtypes, and remained significantly increased for as long as 60 days after ScNT in NF2 neurons. The data clearly shows that *Cdkn1a* RNA levels peak after three to seven days post-injury, before decreasing again. Notably, *Cdkn1a* expression was significantly increased also in the non-neuronal cell types, although to a lesser extent and not as consistently as in neurons. At 24 days post-paclitaxel application as a model for CIPN, *Cdkn1a* expression was only significantly increased in *Sst*^+^ pruriceptors (SST) in this work (log2fc = 1.56, p-adj. < 0.00001).

**Fig 5.**
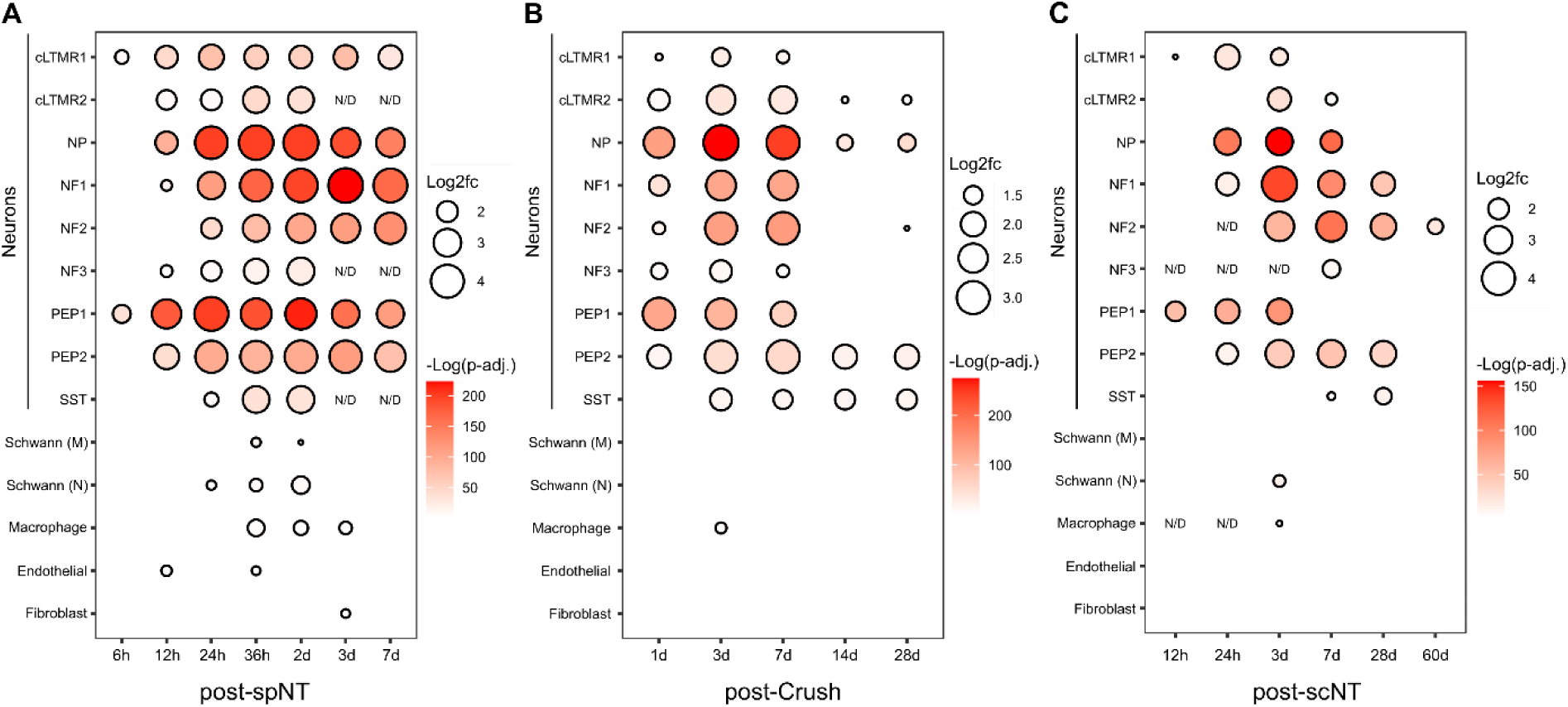
*Cdkn1a* (p21) RNA is increased in all dorsal root ganglion neuronal subtypes following spinal nerve transection, sciatic nerve crush, and sciatic nerve transection in Renthal et al. **(A-C)** Bubble heatmap of significantly increased *Cdkn1a* RNA expression as log2-fold change and -log10(p-adj.) in mouse dorsal root ganglion (DRG) cells following spinal nerve transection (spNT) (A), sciatic nerve crush (Crush) (B), and sciatic nerve transection and ligation (ScNT) (C) when compared to expression in DRG cells from naïve mice, as obtained from Renthal et al. [26]. Differential *Cdkn1a* gene expression is only shown when p-adj. < 0.01 and log2fc > |1| when compared to expression in cells from naïve control mice. Statistical significance was tested in the original manuscript using the QL F-test in edgeR [26]. N/D = not data for these timepoints, because too few cells were available to perform cell type-specific differential gene expression analysis at these individual timepoints.

In the whole study setting, *Cdkn2a* expression was only significantly increased in NF2 neurons at 60 days post-ScNT (log2fc = 1.30, p-adj. < 0.00001).

Overall, this data set substantiates that DRG neurons are the origin of increased *Cdkn1a* RNA levels as observed in the previously described bulk-RNA sequencing and microarray-based studies that investigated neuropathic pain models. Furthermore, it suggests that *Cdkn1a* expression is increased in all identified DRG neuron subtypes, although different subtypes may be differently susceptible or show different rates of neuronal senescence. At last, this study corroborates that *Cdkn2a* is not effectively upregulated following peripheral nerve injury in DRG neurons.

### *Cdkn1a* and *Cdkn2a* RNA is not significantly increased across inflammatory pain models

A prior study suggested that modelling inflammatory pain with CFA leads to senescence of DRG neurons as early as two days following injection [36]. To test whether inducing inflammatory pain is accompanied by increased *Cdkn1a* or *Cdkn2a* RNA in different study settings, we searched for transcriptome studies that applied inflammatory pain models. Parisien et al. performed whole-genome bulk-RNA sequencing with lumbar DRG tissue from mice three days after CFA application, and naive mice as control [25]. This timepoint was selected because von Frey behavioural testing showed that induced hypersensitivity peaked at three days post-CFA [25]. At this day, 839 genes were significantly upregulated in DRG tissue. However, neither *Cdkn1a* nor *Cdkn2a* RNA levels were significantly increased at three days post-CFA in this study [25].

In the previously described study that performed snRNA analysis to investigate gene expression changes in nine DRG neuronal subtypes and non-neuronal cells across four different neuropathic pain models (SpNT, ScNT, Crush, Paclitaxel), the authors additionally analysed gene expression two days following CFA-injection as model for inflammatory pain. Although *Cdkn1a* RNA was significantly increased in DRG cells in their neuropathic pain models, *Cdkna1a* expression was not significantly increased two days post-CFA application [26].

Another model of inflammatory pain is injecting the immune activator zymosan. Strong et al. performed microarray analysis with DRG tissue from rats three days after injection with zymosan in incomplete Freund’s adjuvant beneath the intervertebral foramen onto the L5 DRG to model inflammatory pain, and from sham-treated rats as control [33]. Mechanical hypersensitivity at three days post-injection in zymosan-injected, and lack of hypersensitivity in sham animals, was confirmed by von Frey behavioural testing [33]. In DRG tissue at three days post-zymosan injection, 1625 out of 6831 captured genes showed significantly differential RNA levels (p-adj. < 0.05) including *Cdkn1a* [33]. *Cdkn1a* RNA was increased by 1.37-fold in DRG tissue after zymosan injection when compared to DRG tissue from sham-treated rats (p-adj. = 0.016). *Cdkn2a* RNA levels were not significantly increased at three days post-injection.

Thus, in contrast to changes of *Cdkn1a* RNA in neuropathic pain models, results are less consistent between studies investigating inflammatory pain. It remains unclear whether inducing inflammatory pain in rodents consistently leads to increased *Cdkn1a* in DRG tissue, and whether it may at all lead to increased *Cdkn2a* RNA levels.

### *Cdkn1a* and *Cdkn2a* RNA is not significantly increased in all studies and pain models

We found two additional data sets in which neither *Cdkn1a* nor *Cdkn2a* RNA was significantly increased in rodent DRG tissue upon induction of peripheral-driven pain. In the context of diabetic neuropathy, Athie et al. performed whole-genome bulk-RNA sequencing analysis with DRG tissue from rats four weeks following induction of type 1 diabetes with streptozotocin and control rats [2]. Induction of type 1 diabetes resulted in mechanical hypersensitivity from day 14 on as determined by von Frey behavioural testing and led to 56 significantly expressed genes in DRG cells after four weeks (p-adj. < 0.05). *Cdkn1a* and *Cdkn2a* were not significantly differentially expressed [2]. Kummer et al. performed microarray analysis with DRG tissue isolated from α-galactosidase A-deficient and wildtype mice as a model for Fabry disease-related neuropathic pain [23]. In total, 812 genes were significantly differentially expressed between DRG tissue from α-galactosidase A-deficient and wildtype mice (|Fold change| ≥ 1.2, p-adj. < 0.01)[23]. *Cdkn1a* and *Cdkn2a* were not significantly differentially expressed in DRG tissue in this Fabry disease model.

### *Cdkn1a* RNA is increased by a maximum of 8.26-fold on average in DRG neurons following nerve injury

Due to the varying timepoints, pain models, interventions, sequencing pipelines, and downstream analyses that underly the available data sets, it is very difficult to calculate any mean fold changes of *Cdkn1a* RNA levels that may be expected in future studies when analysing peripheral sensory neuron-specific senescence in chronic pain states. Still, we concluded that *Cdkn1a* RNA levels may peak around day three to seven following nerve injury, and throughout this work, we have analysed three independent studies encompassing eight different differential gene expression data sets on rodent DRG neuron-specific transcriptomes at either day three or seven after lesion-based nerve injury through surgical intervention (SNI, SNT, SpNT, ScNT, and Crush) [3; 17; 26]. Hence, in an attempt to estimate a probable maximum increase in *Cdkn1a* RNA in peripheral sensory neurons upon nerve injury, we calculated an average fold change of *Cdkn1a* RNA levels based on these studies, which is 8.26-fold (±4.50) when compared to *Cdkn1a* RNA levels in control DRG neurons.

## Discussion

Peripheral sensory neuron senescence has recently been associated with increased nociceptive signalling, and suggested as a target for novel pain resolving strategies. By analysing publicly available transcriptomic data sets, we found that the widely-used marker for cellular senescence *Cdkn1a* is consistently upregulated in DRG tissue - specifically in DRG neurons - in lesion-based neuropathic but less consistently in other pain models (Fig. 6).

**Fig 6.**
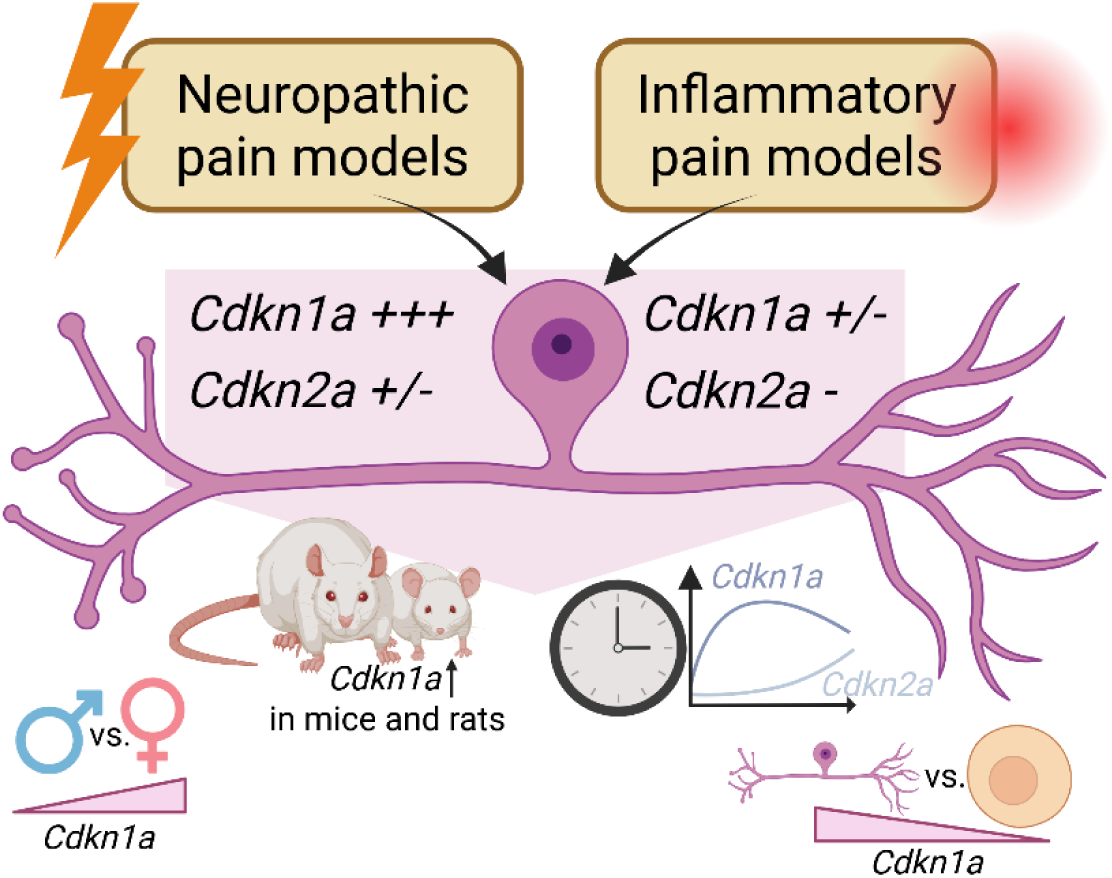
*Cdkn1a* RNA is consistently increased in lesion-based neuropathic pain models. Concluding schematic summarizing key aspects indicated by our analysis of publicly available transcriptomic data sets: *Cdkn1a* RNA may be increased to a higher extent in DRG neurons from females upon nerve injury. *Cdkn1a* RNA is increased in DRG cells from mice and rats in pain conditions. *Cdkn1a* RNA is increased within hours to days following nerve injury while *Cdkn2a* RNA is - if at all - increased only at later timepoints. *Cdkn1a* RNA is increased primarily in DRG neurons and to a lesser extent and less consistently in non-neuronal cells in pain conditions.

We observed that *Cdkn1a* expression is upregulated in DRG neurons in less than a week in lesion-based neuropathic pain models. This is in accordance with other model systems of induced senescence such as drug-induced senescence in cell culture, in which CDKN1A (p21) upregulation can be detected within few hours and steadily increases over the first days [35].

Interestingly, CDKN1A (p21) expression induced in cancer cells by chemotherapy was found to have the capacity to induce both, senescence fate or proliferation [16]. An early increase in p21 did not automatically lead to cellular senescence [16]. Thus, *Cdkn1a* expression at these early timepoints alone may not a guarantee peripheral sensory neuron senescence.

Based on our analysis of data by Renthal et al. [26], we observed that *Cdkn1a* RNA expression levels peak around three to seven days after induction of neuropathic pain, followed by decrease and a final a state in which *Cdkn1a* RNA returns to baseline levels in most DRG neuron subtypes. This is similar to observations for other cell types such as fibroblasts [31]. *Cdkn1a* expression appears to increase during the early phase but subsequently declines once cellular senescence is established. At this stage, senescence-maintaining mechanisms predominate, and SASP develops [14; 31].

Our analysis showed that *Cdkn2a* expression was significantly increased only 30 days post-SNI or 60 days post-ScNT, but not in the studies that performed differential gene expression analysis seven days post-SNI. Hence, increased *Cdkn2a* expression is - if at all - observed at later timepoints in DRG neurons of neuropathic pain models. In a previous study, Donovan et al. likewise observed significantly increased numbers of *Cdkn2a*-positive DRG neurons only at three weeks, but not at seven days post-SNI, when analysing 11-16 weeks-old mice [12]. In contrast, the numbers of *Cdkn1a*-positive neurons were significantly increased already at seven days post-SNI in their study [12]. The here analysed data sets seem to corroborate such a delayed *Cdkn2a* upregulation that occurs after *Cdkn1a* upregulation following SNI in DRG cells, which is a common observation in the context of injury-induced senescence and may be due to the different functions of the two gene products [8; 31; 44]. On the other hand, there is also evidence that distinct cell types and subpopulations either predominantly express high *Cdkn1a* or high *Cdkn2a* levels [44]. Indeed, CDKN2A expression has been previously pointed out as not as reliable as CDKN1A to monitor senescence of central nervous system neurons [14]. Since increased *Cdkn2a* expression was not consistently found in rodent DRG neurons in the here analysed studies, even at later timepoints, this may indicate that *Cdkn2a* expression may not be as meaningful as *Cdkn1a* for peripheral sensory neuron senescence.

We then found evidence that *Cdkn1a* upregulation in DRG tissue in response to nerve injury may be higher in females than in males. Previously, etoposide-induced DNA damage led to higher levels of *Cdkn1a* mRNA in astrocytes from female than from male mice [21]. However, additional evidence for an increased *Cdkn1a* upregulation in female cells is lacking. Nevertheless, we argue that this data set on *Cdkn1a* upregulation in female but not male DRG tissue upon nerve injury calls for more research on sex-specific peripheral sensory neuronal senescence. Moreover, future studies should include female and male animals, which is often not the case (Table 1), and if possible, investigate possible sex-difference.

Given the increase in *Cdkn1a* RNA in DRG neurons across lesion-based neuropathic pain models found in this study, the previously indicated role for Cdkn1a as necessary signal transducer to induce neuronal senescence [20], and the association of senolytic therapy with reduced nocifensive behaviour in animal pain models [12; 36], it would be interesting to test conditional Cdkn1a (p21) knockout mice in such pain models and whether pain chronification is diminished following induction of neuropathic pain. On the other hand, p21 overexpressing (*Cdkn1a*^super^) mice exist, and it would be of great interest to test whether these animals suffer from worse pain phenotypes than wildtype mice based on potentially increased rates of peripheral sensory neuron senescence [37]. Such experimental studies would further corroborate the putative role of Cdkn1a in peripheral-driven pain chronification.

Limitations of our study were the heterogeneity of available transcriptomic studies. Even within single studies like the comprehensive snRNA sequencing work by Renthal et al. [26], timepoints for differential gene expression analysis following the different interventions are not the same. In addition, there are fewer transcriptome studies that have analysed DRG cell gene expression responses in the context of inflammatory pain. This is especially true for later timepoints after CFA or zymosan application. Thus, our picture of *Cdkn1a* and *Cdkn2a* expression as marker for neuronal senescence in the context of peripheral pain is still incomplete. *Cdkn1a* and *Cdkn2a* expression levels are two well-established and widely-used markers for cellular senescence. However, their expression levels alone are not enough to demonstrate and conclude induction of neuronal senescence [18; 34]. Hence, we were careful with our interpretations and think that our study provides a starting point, but that future studies generating transcriptome data sets must analyse additional senescence-related factors. Lastly, we focused on rodent pain models, because these studies are more consistent and better comparable than the few existing studies with human tissue with regard to our purpose of comparing *Cdkn1a* and *Cdkn2a* expression across different study settings.

Overall, we conclude that a senescence-associated increase in *Cdkn1a* expression in DRG neurons is consistently found across rodent lesion-based neuropathic pain models. It seems that increased *Cdkn1a* expression is not as often found in pain models beyond lesion-based nerve injury. The timepoint of analysis, the age and sex of the employed model animals may play important roles when investigating *Cdkn1a* gene expression.

## Acknowledgments

We acknowledge funding from the German Research Foundation (HU 1636/13-1) (PNO, TH) and Köln Fortune (41/2025) (RD, TH).

## Author contributions

Conceptualization: PO, TH

Methodology: PO, RD, JI

Investigation: PO, RD, JI

Visualization: PO

Funding acquisition: TH

Writing - original draft: PO

Writing - review and editing: All authors

## Data availability Statement

The data underlying each figure are available from the corresponding authors upon request.

## Abbreviations

CCI: Chronic constriction injury
CFA: Complete Freund’s adjuvant
CIPN: Chemotherapy-induced peripheral neuropathy
DRG: Dorsal root ganglion
FDR: False discovery rate
GEO: Gene expression omnibus
LCM: Laser capture microdissection
SASP: Senescence-associated secretory phenotype
ScNT: Sciatic nerve transection
SNI: Spared nerve injury
SpNT: Spinal nerve transection

## References

[1] Amorim IS, Kedia S, Kouloulia S, Simbriger K, Gantois I, Jafarnejad SM, Li Y, Kampaite A, Pooters T, Romanò N, Gkogkas CG. Loss of eIF4E Phosphorylation Engenders Depression-like Behaviors via Selective mRNA Translation. J Neurosci 2018;38(8):2118–2133.

[2] Athie MCP, Vieira AS, Teixeira JM, Dos Santos GG, Dias EV, Tambeli CH, Sartori CR, Parada CA. Transcriptome analysis of dorsal root ganglia’s diabetic neuropathy reveals mechanisms involved in pain and regeneration. Life Sci 2018;205:54–62.

[3] Berta T, Perrin FE, Pertin M, Tonello R, Liu YC, Chamessian A, Kato AC, Ji RR, Decosterd I. Gene Expression Profiling of Cutaneous Injured and Non-Injured Nociceptors in SNI Animal Model of Neuropathic Pain. Sci Rep 2017;7(1):9367.

[4] Borsook D, Hargreaves R, Bountra C, Porreca F. Lost but making progress--Where will new analgesic drugs come from? Sci Transl Med 2014;6(249):249sr243.

[5] Breivik H, Collett B, Ventafridda V, Cohen R, Gallacher D. Survey of chronic pain in Europe: prevalence, impact on daily life, and treatment. Eur J Pain 2006;10(4):287–333.

[6] Calls A, Torres-Espin A, Navarro X, Yuste VJ, Udina E, Bruna J. Cisplatin-induced peripheral neuropathy is associated with neuronal senescence-like response. Neuro Oncol 2021;23(1):88–99.

[7] Chaib S, Tchkonia T, Kirkland JL. Cellular senescence and senolytics: the path to the clinic. Nat Med 2022;28(8):1556–1568.

[8] Chandra A, Lagnado AB, Farr JN, Monroe DG, Park S, Hachfeld C, Tchkonia T, Kirkland JL, Khosla S, Passos JF, Pignolo RJ. Targeted Reduction of Senescent Cell Burden Alleviates Focal Radiotherapy-Related Bone Loss. J Bone Miner Res 2020;35(6):1119–1131.

[9] Clough E, Barrett T, Wilhite SE, Ledoux P, Evangelista C, Kim IF, Tomashevsky M, Marshall KA, Phillippy KH, Sherman PM, Lee H, Zhang N, Serova N, Wagner L, Zalunin V, Kochergin A, Soboleva A. NCBI GEO: archive for gene expression and epigenomics data sets: 23-year update. Nucleic Acids Res 2024;52(D1):D138–D144.

[10] Cobos EJ, Nickerson CA, Gao F, Chandran V, Bravo-Caparrós I, González-Cano R, Riva P, Andrews NA, Latremoliere A, Seehus CR, Perazzoli G, Nieto FR, Joller N, Painter MW, Ma CHE, Omura T, Chesler EJ, Geschwind DH, Coppola G, …, Costigan M. Mechanistic Differences in Neuropathic Pain Modalities Revealed by Correlating Behavior with Global Expression Profiling. Cell Rep 2018;22(5):1301–1312.

[11] Dib-Hajj SD, Waxman SG. Translational pain research: Lessons from genetics and genomics. Sci Transl Med 2014;6(249):249sr244.

[12] Donovan LJ, Brewer CL, Bond SF, Laslavic AM, Pena Lopez A, Colman L, Jordan CE, Hansen LH, González OC, Pujari A, de Lecea L, Quarta M, Kauer JA, Tawfik VL. Aging and injury drive neuronal senescence in the dorsal root ganglia. Nat Neurosci 2025;28(5):985–997.

[13] Edgar R, Domrachev M, Lash AE. Gene Expression Omnibus: NCBI gene expression and hybridization array data repository. Nucleic Acids Res 2002;30(1):207–210.

[14] Herdy JR, Mertens J, Gage FH. Neuronal senescence may drive brain aging. Science 2024;384(6703):1404–1406.

[15] Herdy JR, Traxler L, Agarwal RK, Karbacher L, Schlachetzki JCM, Boehnke L, Zangwill D, Galasko D, Glass CK, Mertens J, Gage FH. Increased post-mitotic senescence in aged human neurons is a pathological feature of Alzheimer’s disease. Cell Stem Cell 2022;29(12):1637–1652.e1636.

[16] Hsu CH, Altschuler SJ, Wu LF. Patterns of Early p21 Dynamics Determine Proliferation-Senescence Cell Fate after Chemotherapy. Cell 2019;178(2):361–373.e312.

[17] Hu G, Huang K, Hu Y, Du G, Xue Z, Zhu X, Fan G. Single-cell RNA-seq reveals distinct injury responses in different types of DRG sensory neurons. Sci Rep 2016;6:31851.

[18] Hudson HR, Riessland M, Orr ME. Defining and characterizing neuronal senescence, ‘neurescence’, as G. Trends Neurosci 2024;47(12):971–984.

[19] Johannes CB, Le TK, Zhou X, Johnston JA, Dworkin RH. The prevalence of chronic pain in United States adults: results of an Internet-based survey. J Pain 2010;11(11):1230–1239.

[20] Jurk D, Wang C, Miwa S, Maddick M, Korolchuk V, Tsolou A, Gonos ES, Thrasivoulou C, Saffrey MJ, Cameron K, von Zglinicki T. Postmitotic neurons develop a p21-dependent senescence-like phenotype driven by a DNA damage response. Aging Cell 2012;11(6):996–1004.

[21] Kfoury N, Sun T, Yu K, Rockwell N, Tinkum KL, Qi Z, Warrington NM, McDonald P, Roy A, Weir SJ, Mohila CA, Deneen B, Rubin JB. Cooperative p16 and p21 action protects female astrocytes from transformation. Acta Neuropathol Commun 2018;6(1):12.

[22] Kharchenko PV, Silberstein L, Scadden DT. Bayesian approach to single-cell differential expression analysis. Nat Methods 2014;11(7):740–742.

[23] Kummer KK, Kalpachidou T, Kress M, Langeslag M. Signatures of Altered Gene Expression in Dorsal Root Ganglia of a Fabry Disease Mouse Model. Front Mol Neurosci 2017;10:449.

[24] Ng M, Hazrati LN. Evidence of sex differences in cellular senescence. Neurobiol Aging 2022;120:88–104.

[25] Parisien M, Samoshkin A, Tansley SN, Piltonen MH, Martin LJ, El-Hachem N, Dagostino C, Allegri M, Mogil JS, Khoutorsky A, Diatchenko L. Genetic pathway analysis reveals a major role for extracellular matrix organization in inflammatory and neuropathic pain. Pain 2019;160(4):932–944.

[26] Renthal W, Tochitsky I, Yang L, Cheng YC, Li E, Kawaguchi R, Geschwind DH, Woolf CJ. Transcriptional Reprogramming of Distinct Peripheral Sensory Neuron Subtypes after Axonal Injury. Neuron 2020;108(1):128–144.e129.

[27] Rice ASC, Smith BH, Blyth FM. Pain and the global burden of disease. Pain 2016;157(4):791–796.

[28] Riessland M, Kolisnyk B, Kim TW, Cheng J, Ni J, Pearson JA, Park EJ, Dam K, Acehan D, Ramos-Espiritu LS, Wang W, Zhang J, Shim JW, Ciceri G, Brichta L, Studer L, Greengard P. Loss of SATB1 Induces p21-Dependent Cellular Senescence in Post-mitotic Dopaminergic Neurons. Cell Stem Cell 2019;25(4):514–530.e518.

[29] Ritchie ME, Phipson B, Wu D, Hu Y, Law CW, Shi W, Smyth GK. limma powers differential expression analyses for RNA-sequencing and microarray studies. Nucleic Acids Res 2015;43(7):e47.

[30] Starobova H, Mueller A, Deuis JR, Carter DA, Vetter I. Inflammatory and Neuropathic Gene Expression Signatures of Chemotherapy-Induced Neuropathy Induced by Vincristine, Cisplatin, and Oxaliplatin in C57BL/6J Mice. J Pain 2020;21(1-2):182–194.

[31] Stein GH, Drullinger LF, Soulard A, Dulić V. Differential roles for cyclin-dependent kinase inhibitors p21 and p16 in the mechanisms of senescence and differentiation in human fibroblasts. Mol Cell Biol 1999;19(3):2109–2117.

[32] Stephens KE, Zhou W, Ji Z, Chen Z, He S, Ji H, Guan Y, Taverna SD. Sex differences in gene regulation in the dorsal root ganglion after nerve injury. BMC Genomics 2019;20(1):147.

[33] Strong JA, Xie W, Coyle DE, Zhang JM. Microarray analysis of rat sensory ganglia after local inflammation implicates novel cytokines in pain. PLoS One 2012;7(7):e40779.

[34] Suryadevara V, Hudgins AD, Rajesh A, Pappalardo A, Karpova A, Dey AK, Hertzel A, Agudelo A, Rocha A, Soygur B, Schilling B, Carver CM, Aguayo-Mazzucato C, Baker DJ, Bernlohr DA, Jurk D, Mangarova DB, Quardokus EM, Enninga EAL, …, Neretti N. SenNet recommendations for detecting senescent cells in different tissues. Nat Rev Mol Cell Biol 2024;25(12):1001–1023.

[35] te Poele RH, Okorokov AL, Jardine L, Cummings J, Joel SP. DNA damage is able to induce senescence in tumor cells in vitro and in vivo. Cancer Res 2002;62(6):1876–1883.

[36] Techameena P, Feng X, Zhang K, Hadjab S. The single-cell transcriptomic atlas iPain identifies senescence of nociceptors as a therapeutical target for chronic pain treatment. Nat Commun 2024;15(1):8585.

[37] Torgovnick A, Heger JM, Liaki V, Isensee J, Schmitt A, Knittel G, Riabinska A, Beleggia F, Laurien L, Leeser U, Jüngst C, Siedek F, Vogel W, Klümper N, Nolte H, Wittersheim M, Tharun L, Castiglione R, Krüger M, …, Reinhardt HC. The Cdkn1a. Cell Rep 2018;25(4):1027–1039.e1026.

[38] Tsang A, Von Korff M, Lee S, Alonso J, Karam E, Angermeyer MC, Borges GL, Bromet EJ, Demytteneare K, de Girolamo G, de Graaf R, Gureje O, Lepine JP, Haro JM, Levinson D, Oakley Browne MA, Posada-Villa J, Seedat S, Watanabe M. Common chronic pain conditions in developed and developing countries: gender and age differences and comorbidity with depression-anxiety disorders. J Pain 2008;9(10):883–891.

[39] Uttam S, Wong C, Amorim IS, Jafarnejad SM, Tansley SN, Yang J, Prager-Khoutorsky M, Mogil JS, Gkogkas CG, Khoutorsky A. Translational profiling of dorsal root ganglia and spinal cord in a mouse model of neuropathic pain. Neurobiol Pain 2018;4:35–44.

[40] Vega-Avelaira D, Géranton SM, Fitzgerald M. Differential regulation of immune responses and macrophage/neuron interactions in the dorsal root ganglion in young and adult rats following nerve injury. Mol Pain 2009;5:70.

[41] Wickham H. ggplot2 : Elegant Graphics for Data Analysis. Use R!,. Cham: Springer International Publishing : Imprint: Springer,, 2016. pp. 1 online resource (XVI, 260 pages 232 illustrations, 140 illustrations in color.

[42] Woolf CJ. Overcoming obstacles to developing new analgesics. Nat Med 2010;16(11):1241–1247.

[43] Xiao Y, Hsiao TH, Suresh U, Chen HI, Wu X, Wolf SE, Chen Y. A novel significance score for gene selection and ranking. Bioinformatics 2014;30(6):801–807.

[44] Yan J, Chen S, Yi Z, Zhao R, Zhu J, Ding S, Wu J. The role of p21 in cellular senescence and aging-related diseases. Mol Cells 2024;47(11):100113.

